# A hemifused complex is the hub in a network of pathways to membrane fusion

**DOI:** 10.1101/2021.06.20.449175

**Authors:** Jason M. Warner, Dong An, Benjamin S. Stratton, Ben O’Shaughnessy

## Abstract

Membrane fusion is a critical step for many essential processes, from neurotransmission to fertilization. For over 40 years protein-free fusion driven by calcium or other cationic species has provided a simplified model of biological fusion, but the mechanisms remain poorly understood. Cation-mediated membrane fusion and permeation are essential in their own right to drug delivery strategies based on cell-penetrating peptides or cation-bearing lipid nanoparticles. Experimental studies suggest calcium drives anionic membranes to a hemifused intermediate which constitutes a hub in a network of pathways, but the pathway selection mechanism is unknown. Here we develop a mathematical model that identifies the network hub as a highly dynamical hemifusion complex. Multivalent cations drive expansion of this high tension hemifusion interface between interacting vesicles during a brief transient. The fate of this interface determines the outcome, either fusion, dead-end hemifusion or vesicle lysis. The model reproduces the unexplained finding that calcium-driven fusion of vesicles with planar membranes typically stalls at hemifusion, and we show the equilibrated hemifused state is a novel lens-shaped complex. Thus, membrane fusion kinetics follow a stochastic trajectory within a network of pathways, with outcome weightings set by a hemifused complex intermediate.

**Significance:** Cells use multicomponent machineries to fuse membranes for neurotransmitter and hormone release and other fundamental processes. Protein-free fusion using calcium or other multivalent cationic fusogens has long been studied as a simplifying model. Cation-mediated membrane fusion or permeation are key events for a number of current drug delivery strategies. However, the mechanisms determining outcomes are unknown. Here we develop a mathematical model that identifies a dynamic hemifusion complex as the decision hub that stochastically sets the outcome in a network of pathways. Cations transiently grow a high tension hemifusion interface between membrane-enclosed compartments, whose fate governs whether fusion, dead-end hemifusion or vesicle lysis occurs. The model provides a systematic framework to predict outcomes of cationic fusogen-mediated interactions between membrane-enclosed compartments.

## Introduction

Membrane fusion is essential for exocytosis, intracellular trafficking, fertilization and other processes vital for living organisms (1, 2). Fusion in cells is regulated by multicomponent cellular machineries, many aspects of which remain poorly understood (3, 4). For over 40 years protein-free fusion has served as a simplified model of biological fusion, retaining the complexities of phospholipid membranes but with simplified fusogens, most commonly calcium or other divalent cations (5-11). Protein-free systems could help elucidate the pathways to biological fusion and the biophysical properties of lipid bilayers that cellular fusion machineries must contend with over many scales, from ∼50-nm synaptic vesicles to cortical granules with sizes from ∼ 100 nm to several microns (12-17).

Cation-mediated fusion and membrane permeation also find use in a number of biotechnologies for delivery of drugs and other cargoes. Cationic cell-penetrating peptides (CPPs) with < 30 amino acids have been widely studied as agents of drug delivery for cancer, central nervous system diseases and others (18, 19), either covalently bonded to small cargo molecules or physically adhered to negatively charged cargoes such as DNA or RNA (18, 19). Examples of CPPs include a polybasic sequence from TAT, a transcription activator protein of HIV (20), and the model CPP nona-arginine (R9) (21). How cargoes are delivered is not established, but CPPs increase membrane permeability (22-25) and R9 was shown to fuse large vesicles (22). Similarly, antimicrobial peptides are naturally occurring and synthetically mimicked antibiotics, often positively charged, that permeabilize target membranes (26).

By far the most studied cationic fusogen is calcium. Historically calcium was proposed to be used by cells as a fusogen (27). Early in vitro studies in bulk suspension (5-8, 28) and later single event methods tracking Ca^2+^-mediated fusion of vesicles with planar bilayers (black lipid membranes) (29-31) identified several calcium-driven processes on the pathway to fusion. Adhesion of membranes was followed by hemifusion, a preliminary step fusing the outer monolayers only. The hemifused state was the final outcome (“dead-end hemifusion”), or hemifusion evolved into either full fusion or vesicle lysis, the rupture of a non-contacting membrane surface.

These discoveries suggest hemifusion is the critical intermediate on the fusion pathway. Indeed, in biological contexts hemifused intermediates are seen in live cells (16) and *in vitro* driven by SNARE proteins and other cellular fusion machinery components (32, 33), while hemifusion is observed in molecular dynamics simulations (34, 35). Explicit support for hemifusion as the intermediate that sets the fusion outcome was provided by a study in which divalent cations adhered and hemifused giant unilamellar vesicles (GUVs) (11). At high calcium concentrations (6 mM Ca^2+^) the hemifused connections grew within seconds into micron sized hemifusion diaphragms (HDs) which ruptured to give fusion, or else vesicle lysis occurred. Lower concentrations produced dead-end hemifusion, with equilibrated HDs. A mathematical model (36) reproducing the observed growth rates and HD sizes explained the HD expansion as driven by the powerful tendency of Ca^2+^ to contract anionic lipid monolayers and bilayers (6, 7, 37, 38) and dramatically increase the membrane tension as a result.

The findings of ref. (11) suggest the pathway to membrane fusion is hemifusion, followed by rupture of an expanding HD. Consistent with this pathway, Ca^2+^ typically can only achieve dead-end hemifusion of GUVs with planar membranes (PMs), but fusion can be subsequently triggered by osmotic boosting of membrane tension or application of spontaneous curvature lysolipids to the distal PM monolayer (29-31). This suggests positive curvature pores expand to rupture the HD, but only if pore curvature energetics permit or if the membrane tension is sufficient. Negative curvature phospholipids such as PE have the opposite effect, as Ca^2+^ fuses or lyses pure PS GUVs, but lysis is abolished when PE is present (9, 10).

In summary, multivalent cations catalyse membrane fusion via a network of pathways passing through the hemifused state, but the pathway selection mechanism is unknown. Here we develop a mathematical analysis that predicts the pathway weightings and quantitatively reproduces the principal experimental findings. We show that, following nucleation, the growing HD has very high tension and may rupture (fusion outcome) or survive the episode and equilibrate (dead-end hemifusion). A third possibility is vesicle membrane rupture (lysis outcome). The model reproduces the unexplained finding that, unless assisted by osmotic pressure or positive curvature lipids, fusion of vesicles with planar membranes typically stalls at hemifusion, and we show that this hemifused state is a novel lens-shaped complex. Thus, membrane fusion kinetics follow a stochastic trajectory within a network of pathways, with outcome weightings set by a hemifused complex intermediate which determines the dependence on fusogen concentration, vesicle size, lipid composition and geometry.

## Results

### Model

We consider a pair of anionic lipid vesicles in the presence of Ca^2+^ (Fig. 1A). Calcium strongly adheres negatively charged or zwitterionic lipid vesicles, with adhesion energy *W* per unit area dependent on cation concentration and lipid composition (39, 40). Experimentally, the strong adhesion catalyzes hemifusion of GUVs after ∼1-10 s (30, 31, 41) and nucleates a HD (11).

**Figure 1.**
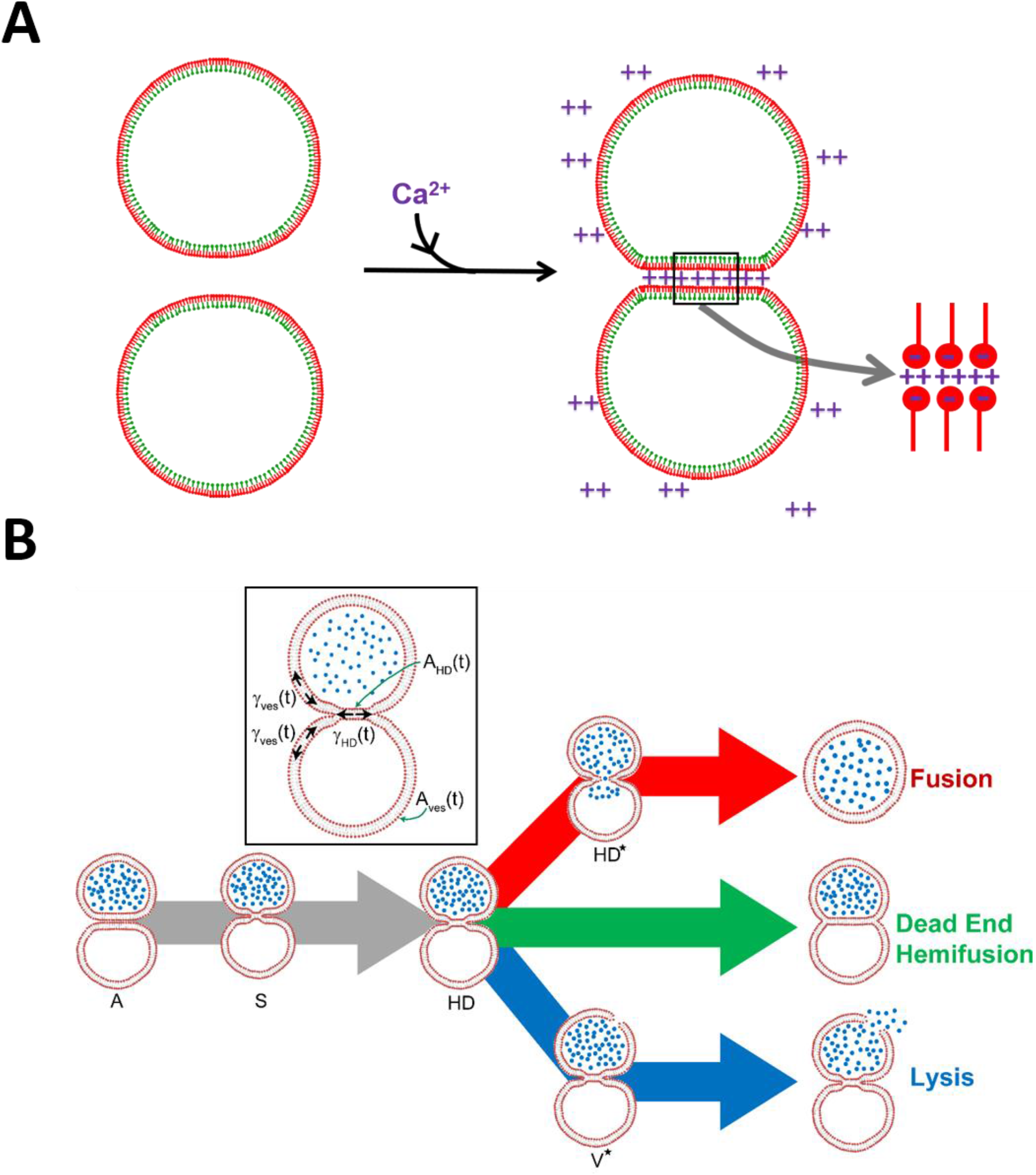
(A) Ca^2+^ increases membrane tension and adheres negatively charged vesicles. Ca^2+^ interacts with anionic or zwitterionic lipids in the outer leaflet of the vesicle membrane to contract the vesicle membrane by a factor 𝜖 to increase tension. A second effect is to adhere the vesicles, with adhesion energy *W* per unit area. (B) Network of pathways to Ca^2+^-mediated membrane fusion. Calcium and other divalent cations adhere phospholipid bilayer vesicle membranes (state **A**) and provoke hemifusion, fusion of the outer phospholipid monolayers only. The initial hemifusion connection is thought to be a minimal stalk (state **S**) that is metastable and yields to an expanding hemifusion diaphragm, HD (state **HD**). HD expansion is driven by high calcium-induced membrane tension and outer monolayer contraction. The HD bilayer tension is greatest and may generate a pore (state **HD\***) that could reseal or dilate and rupture the HD (fusion outcome); or the vesicle membrane could nucleate a pore (state **V\***) that dilates and causes rupture (lysis outcome); or the HD may survive the high tension transient unscathed, expanding to full equilibrium (dead-end hemifusion outcome). Inset: The transient hemifused state (**HD**) selects the pathway. The HD tension γ_*HD*_*(t)* is greatest, as it balances two vesicle tensions γ_*ves*_(t), but its area is least, *A*_*HD*_*(t) < A*_*ves*_*(t)*. These effects compete to set the ratio of fusion to lysis. Since tension rapidly decays as the HD expands, the HD may escape to low tension equilibrium (dead-end hemifusion).

Once nucleated, the HD will rapidly expand due to two forces. First, Ca^2+^ boosts membrane tension (e.g. 8 mN/m for DOPS monolayers with 2 mM Mg^2+^ (6)), since Ca^2+^ tends to contract membranes by factors 𝜖_cat_ ≈ 5 −9% (6, 7, 37, 38). This increased membrane tension drives HD growth, since tension favors smaller areas and a larger HD decreases the total area of the inner and outer leaflets in the hemifused complex (36) (Fig. 1A). Second, Ca^2+^ selectively contracts the outer bilayer leaflets, also favoring HD growth.

In the hemifused complex, we will refer to the membrane not belonging to the HD as the vesicle membrane. To grow the HD, the inner and outer leaflets of these vesicle membranes must slide relative to each other (Fig. S1). This produces a difference in their leaflet lipid densities Δ*ρ*, which builds up interleaflet tension in that region,

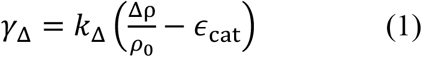

where the interleaflet modulus has typical values *k*_Δ_∼ 20mN/m (42), and *ρ*_0_ is the initial density. Thus, calcium drives sliding of the vesicle membrane leaflets (and hence HD growth) since it favors a non-zero relative density difference, 𝜖_cat_.

The evolution of the leaflet density difference in the vesicle membrane region is set by the balance between the force due to interleaflet tension and the interleaflet drag force, *λ*Δv= −∇*γ*_Δ_, where Δv is the difference in leaflet velocies and *λ* the interleaflet friction coefficient. Using the continuity equation *𝜕*Δ*ρ*/*𝜕t* = −*ρ*_0_∇ ⋅ (Δv) yields the density evolution dynamics

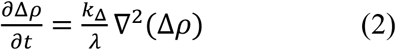

These are to be solved subject to eq. (1) at the vesicle-HD boundary, relating Δ*ρ* to the interleaflet tension *γ*_Δ_. To convert this to a condition invoving the vesicle tension, we use the fundamental relation at this location between the HD, vesicle and interleaflet tensions, *γ*_*HD*_ = *γ*_*ves*_*+*2*γ*_Δ_ (36). Further, the force balance at that location is *γ*_*HD*_ ≈ 2 *γ*_ves_, valid for small contact angle *θ* (Fig. S1), yielding *γ*_Δ_ ≈ *γ*_ves_/2. Using this in eq. (1) gives the boundary condition\

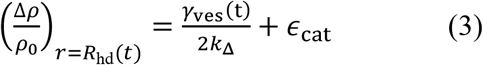

where *R*_*hd*_(*t*) is the HD radius (the origin of coordinates is the HD center, *r* = 0). Finally, the HD outer edge velocity *dR*_*hd*_/*dt* equals Δvat that location, equal to −∇*γ*_Δ_/*λ*. From eq. (1) this gives

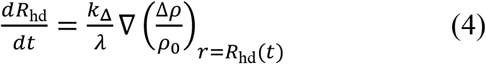

In summary, our procedure is as follows (36). We solve the density difference dynamics in the vesicle membrane region, eq. (2), subject to the moving boundary condition at the moving HD edge, eq. (3). Then eq. (4) gives the HD edge velocity, allowing continuous update of the boundary condition location. The solution yields the increasing HD size and area, *A*_HD_(*t*).

The HD expansion is driven by the vesicle membrane tension *γ*_ves_(*t*) appearing in the boundary condition, eq. (3). Initially very high due to calcium, the tension decays with time due to three principal effects, all accounted for in our model (see SI text). (i) As the HD grows, the total leaflet area (inner plus outer) decreases. (ii) Slow water leakage on sec timescales decreases the vesicle volumes. (iii) The tension decrease due to (i) and (ii) is buffered somewhat by vesicle-vesicle adhesion, which flattens the vesicles and boosts tension.

### Membrane rupture kinetics

The above analysis yields the time-dependent areas and tensions of the HD and vesicle membranes. For high calcium concentrations, the tensions are sufficient to provoke rupture (see below). Three outcomes are then possible. (1) Fusion, if the HD ruptures. (2) Vesicle lysis (contents leakage), if the vesicle membrane ruptures first. (3) Dead end hemifusion, if the HD reaches equilibrium and no rupture occurs.

In ref. (43) higher tensions were shown experimentally to rupture membranes more rapidly. Adapting the mathematical model of tension-mediated rupture from that study, we arrive at the following kinetic scheme

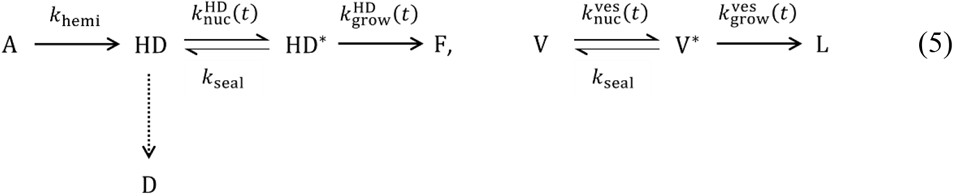

Here **A** is the initial tightly adhered vesicle state; **HD** and **HD\***are states with an expanding HD, with or without a nucleated pore in the HD, respectively; **F** is the fused state following pore growth and HD rupture (Fig. 1B). The second kinetic scheme describes simultaneous processes in the non-HD vesicle membranes: **V** and **V^∗^** denote vesicle membrane states with or without a pore, respectively, and **L** the lysis state. Finally, **D** denotes the dead end hemifused state.

As HD nucleation times have not been documented to our knowledge, for simplicity we assume this first step **A → HD** is instantaneous (for the effects of finite HD nucleation times, see

*nuc*

SI Fig. S4). The pore nucleation rate 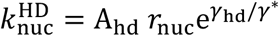, and the rate constant for pore growth and HD lysis 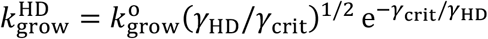, depend on the time-dependent HD area and tension. Here *γ*_crit_ ≡ *𝜋𝜏*^2^/*k*_B_*T* where 𝜏 is the pore line tension (see SI text, eqs. S17, S18) (43, 44). Analogous expressions for rate constants apply to the vesicle membrane lysis kinetics, but with the time-dependent vesicle area and tension.

### Method of calculation

Table 1 lists the parameter values for each experimental system to which we applied our model. We determined the Ca^2+-^-induced contraction factor 𝜖_cat_ and initial membrane tension from a composition weighted average of measured values for pure lipid species (6, 7, 37, 38). Pore line tension 𝜏 was computed from spontaneous curvatures of pure lipid species, and we extended an earlier model (45) to include curvature dependent lipid partitioning effects (see SI text). The line tension 𝜏 is a critical factor, as larger 𝜏 leads to slower pore growth rates 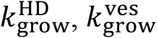, and slower membrane rupture. Adhesion energies *W* were inferred from experimentally reported contact angles. With these parameters, we evolved the HD and vesicle areas and tensions as described above, and used these in the coupled differential equations representing the kinetic scheme of eq. (5). The solutions yielded the fraction of events whose final outcome was fusion, dead-end hemifusion or lysis. For details, see SI text.

**Table 1.** Model parameters. Each line corresponds to one experiment to which the model was applied. Line tension τ and Ca^2+^-induced initial tensions *γ*^0^ and cation contraction factors ϵcat were calculated from single species properties and the lipid composition (see SI text). Adhesion energies *W* were estimated from the observed vesicle-substrate (11) or vesicle-vesicle (9, 10) contact angles (see *SI Text*). Pore kinetics rate parameters are taken as reported in ref. (43), based on rupture measurements of pure DOPC vesicles: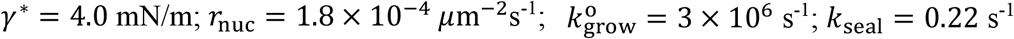.. Lipid abbreviations: PC: phosphocholine; PS: phosphoserine; PE: phosphoethanolamine; LPC: lysophosphatidylcholine; Aso: asolectin; Erg: ergosterol.

| System | Ref. | Experimental Conditions |  |  | Model Parameter Values |  |  |  |  |
| --- | --- | --- | --- | --- | --- | --- | --- | --- | --- |
| | | $R_{\text{ves}}$<br>($\mu\text{m}$) | [Cation] | Composition | $\gamma^0$<br>(mN/m) | $\epsilon_{\text{cat}}$<br>(%) | $W$<br>(mN/m) | $\tau$<br>(pN) | $\gamma_{\text{crit}}$<br>(mN/m) |
| GUV-GUV | (11) | 9 | 6 mM $\text{Ca}^{2+}$ | 60 PC/20 PS/20 PE | 8.6 | 6.3 | 1.1 | 12.8 | 125 |
| GUV-GUV | (11) | 9 | 2 mM $\text{Mg}^{2+}$ | 60 PC/20 PS/20 PE | 7.2 | 5 | 1.1 | 12.8 | 125 |
| GUV-GUV | (9) | 5 | 5 mM $\text{Ca}^{2+}$ | 100 PS | 9.4 | 7 | 5.3 | 12.6 | 122 |
| GUV-GUV | (10) | 5 | 5 mM $\text{Ca}^{2+}$ | 25 PS/75 PE | 8.8 | 6.7 | 5.3 | 16.8 | 215 |
| GUV-PM | (30) | 5 | 20 mM $\text{Ca}^{2+}$ | 80 Aso/20 LPC | 12.2 | 9.2 | - | 7 | 40 |
| GUV-PM | (30) | 5 | 20 mM $\text{Ca}^{2+}$ | 100 Aso | 12.2 | 9.2 | - | 12 | 114 |
| GUV-PM | (31) | 5 | 20 mM $\text{Ca}^{2+}$ | 88 Aso/12 Erg | 10 | 7.5 | - | 13 | 127 |

### Fusion is the domant outcome at intermediate cation concentrations

The model predicts that *Ca*^2*+*^and other cationic fusogens drive membranes along a network of pathways whose hub is the hemifused state (Fig. 1B). How does the outcome distribution depend on the concentration of fusogenic divalent cations? We applied the model to the conditions of the GUV-GUV fusion experiments of ref. (11), accounting for the vesicle dimensions, phospholipid composition and adhesion energies (Table 1).

We calculated outcome distributions over a range of [Ca^2+^] values. Three regimes emerged (Fig. 2A). (1) *Low concentrations*, [Ca^2+^] ≲ 2mM. Following HD nucleation, membrane tension drives HD expansion, but the HD membrane tension is always below the rupture threshold (see below). With high probability the HD therefore expands without rupture to its final equilibrium low-tension state, i.e. dead-end hemifusion results with occasional fusion events. As the vesicle membrane tension is even lower, lysis is improbable. (2) *Intermediate concentrations*, 2 mM ≲ [Ca^2+^] ≲ 10 mM. The initial HD tension now exceeds the rupture threshold, but the vesicle tension does not. Thus HD rupture (fusion outcome) is far more likely than vesicle rupture (lysis outcome). Dead-end hemifusion is rare. (3) *High concentrations*, [Ca^2+^] ≳ 10 mM. All initial tensions exceed the rupture thresholds, but the vesicle has far greater area than does the HD, i.e. many more membrane locations for nucleation of pores that are the precursors of rupture. Thus, lysis is most probable.

**Figure 2.**
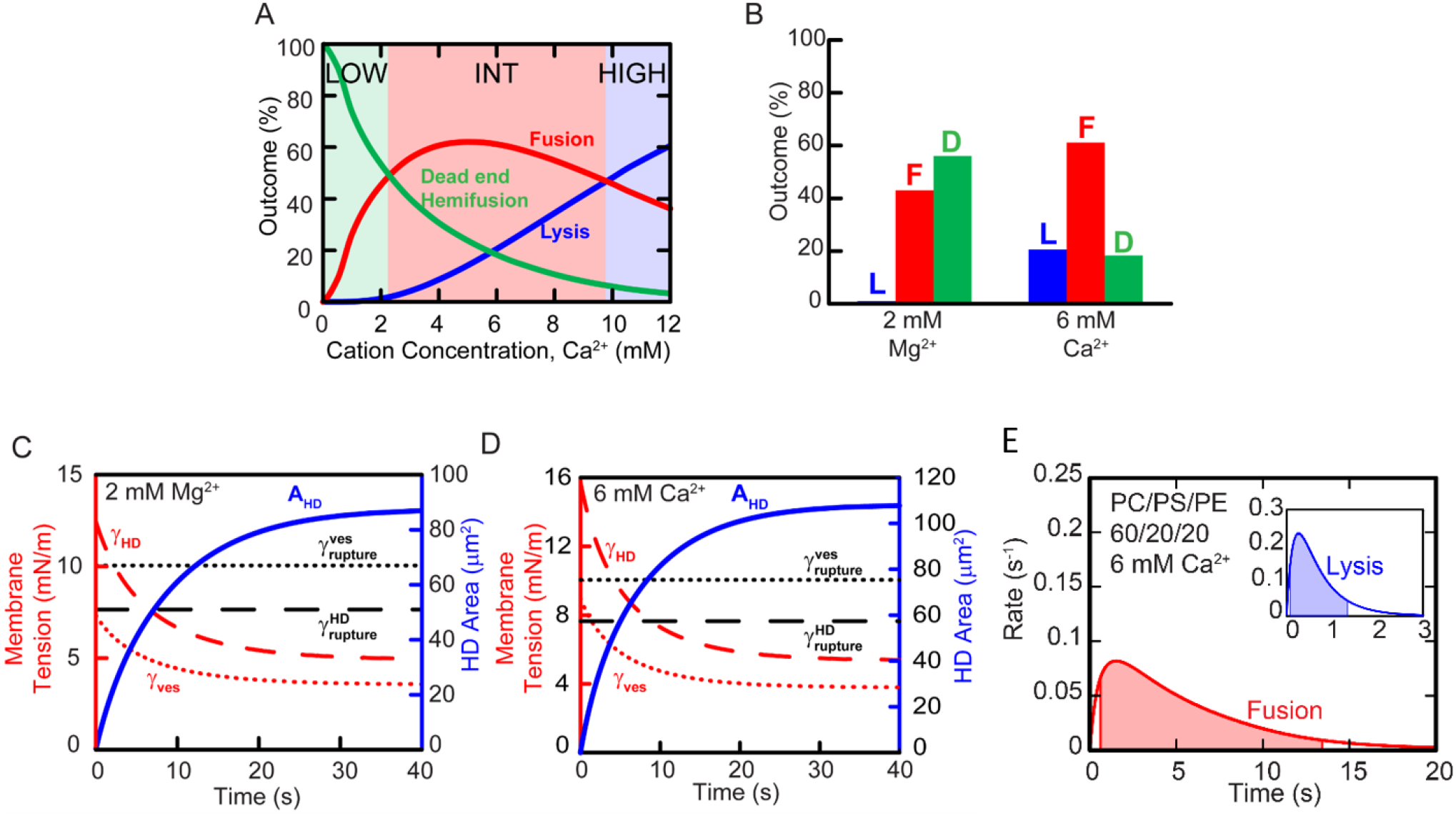
Calcium concentration governs outcome distribution in Ca^2+^-mediated GUV-GUV fusion. Model predictions, conditions as in experiments of ref. (11) (see Table 1 for parameters). GUVs are made of phospholipid with PS/PE/PC = 1/1/3. (A) Predicted outcome distributions versus calcium concentration. In the low concentration regime dead-end hemifusion is most probable; fusion is maximized in the intermediate regime; lysis dominates at high concentrations. (B) Predicted outcome distributions for the two specific cation concentrations used in ref. (11). D, dead-end hemifusion; L, Lysis; F, fusion. (C, D) Predicted time evolution of HD area and tension and vesicle tension at 2 mM Mg^2+^ (C) and 6 mM Ca^2+^(D). HD growth decays the vesicle and HD tensions to below their respective rupture thresholds. More fusion occurs at the higher concentration because the HD tension exceeds its rupture threshold for longer. Calcium only interacts with the PS component in the non-HD region to switch its spontaneous curvature to a negative pore-hating value. Compared with HDs, non-HD membranes are less likely to rupture. (E) Model predicted rates of Ca^2+^-mediated GUV-GUV fusion and lysis versus time for the conditions of the experiments of refs (11).

### Fusion occurs during a limited time window when HD tension is high

Next we compared our model to the findings of ref. (11), where Ca^2+^-mediated interactions between GUVs were studied at two cation concentrations, and growing HDs were visualized from the instant of nucleation (see Table 1 for parameters). At the lower concentration, 2 mM Mg^2+^, the model predicts these experiments lie near the low/intermediate regime boundary (Fig. 2A), with similar outcome probabilities for dead-end hemifusion (56%) and fusion (43%) but almost no lysis (Fig. 2B). This is consistent with ref. (11), where stable dead-end hemifusion events were reported for these conditions (no attempt was made to record fusion or lysis events). For 6 mM Ca^2+^, conditions lie deep within the intermediate regime and the model predicts mainly fusion (61%), with roughly equal probabilities of dead-end hemifusion (18%) or lysis (21%) (Fig. 2B). This outcome distribution is consistent with the reported 2 fusion events and 1 lysis event (11), though the small number of observations precludes firm conclusions.

Fusion can only occur in a limited window of time as the HD grows and its tension decays. At the lower 2 mM cation concentration the HD tension initially exceeds the rupture threshold *γ*_rupture_, where pore growth and resealing rates are equal (*k*_grow_ = *k*_seal_, eq. (5) and *SI text*) (Fig. 2C). According to the kinetics of eq. (1), even a single nucleated pore will likely grow and rupture a membrane above this tension. Initially the HD area is too small to grow pores, but during a ∼ 7s window the HD tension remains above the rupture threshold as the area increases, yielding a net ∼ 50% HD rupture probability (fusion outcome). Should the HD survive this episode, its tension decays below threshold so rupture is no longer possible (dead-end hemifusion). Almost no lysis occurs as the vesicle membrane tension is always below rupture threshold.

At the high 6 mM Ca^2+^ concentration, the initial HD tension is higher, giving a greater net fusion probability during the super-threshold window (Fig. 2D). Since the vesicle tension is higher, and due to the large vesicle area, lysis is now a significant outcome. The distribution of fusion times peaks later and is broader than the lysis time distribution (Fig. 2E). This is because the initial lysis rate is high due to the large vesicle membrane area, but its tension rapidly drops below the rupture threshold so that lysis ceases, whereas the HD needs time to grow sufficiently before significant pore nucleation occurs, but then maintains a super-rupture tension for ∼ 10 s. The predicted timescales are consistent with the reported times of two fusion events (0.75 and 1.75 s) and one lysis event (1.5 s) (11).

### Negative curvature lipids suppress vesicle lysis by lowering pore enlargement rates

While the fusogen concentration sets the driving force for fusion, the phospholipid bilayer composition sets the susceptibility to that force. To study lipid composition effects we applied the model to the experiments of refs. (9, 10) where fusion of ∼10 μm GUVs was studied in bulk suspension, with varying amounts of the phospholipids PS and PE which have positive and negative spontaneous curvatures, respectively. PS/PE compositions from pure PS to 75% PE were studied. We assumed 5 mM Ca^2+^. A subtlety is that in the presence of Ca^2+^ the anionic lipid PS is thought to develop negative spontaneous curvature (Table S1).

For pure PS vesicles, the model predicts dead-end hemifusion is very rare (3%), with substantial probabilities for fusion (37%) and vesicle lysis (60%) (Fig. 3A). Now the positive spontaneous curvature of PS gives a low rupture threshold tension, and on addition of Ca^2+^ both vesicle and HD tensions are super-threshold (Fig. 3C). Thus one might expect exclusively lysis, since the vesicle has far greater area. However, because the PS curvature switches to a negative pore-hating value in the presence of Ca^2+^, the vesicle has a much higher rupture threshold than the internal HD (Fig. 3C). This mechanism protects the vesicle from rupture, allowing significant fusion (Fig. 3E). These predictions agree qualitatively with the experimental outcomes (∼60% fusion, ∼40% lysis, no hemifusion).

**Figure 3.**
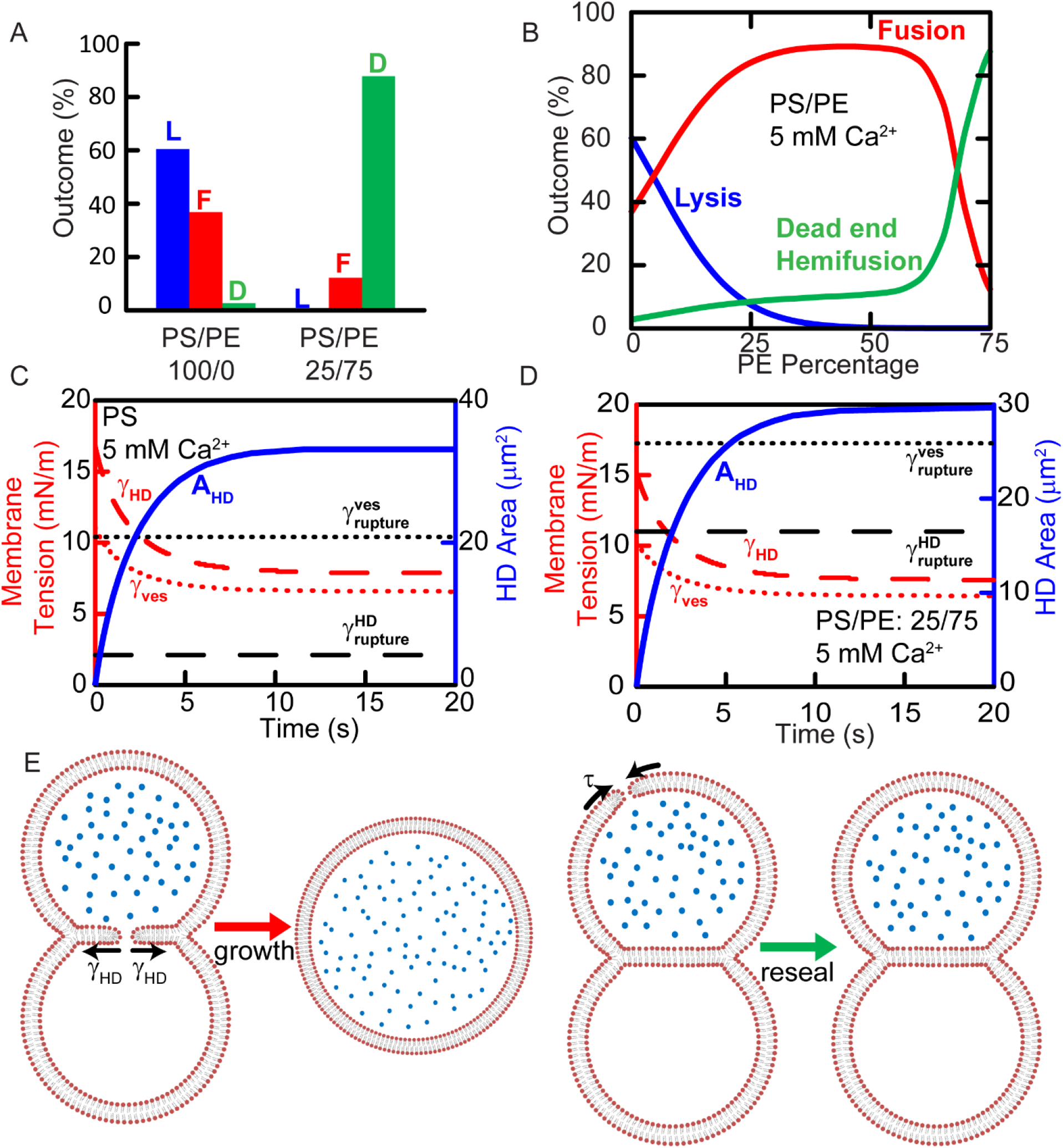
Lipid composition regulates outcome distribution in Ca^2+^-mediated GUV-GUV fusion. GUVs are made of PS and PE with different PE fractions. Model predictions, conditions as in experiments of refs. (9, 10) (see Table 1 for parameters). The negative curvature lipid PE disfavors pore formation and suppresses fusion. (A) Predicted outcome distributions for the compositions of ref. (9) (pure PS) and ref. (10) (75% PE, 25% PS) in the presence of 5 mM Ca^2+^. (B) Predicted outcome distributions versus PS/PE composition in the presence of 5 mM Ca^2+^. (C, D) Predicted time evolution of HD area, vesicle and HD tension for pure PS (C) and 75% PE, 25% PS (D). (E) Mechanism that protects pure PS vesicles from lysis. As PS has positive spontaneous curvature nucleated pores have low line tension and will likely be expanded by the HD membrane tension *γ*_*HD*_ (left). By contrast the pore line tension τ in the vesicle membrane is large, as PS is thought to have negative spontaneous curvature in the presence of Ca^2+^, and pores close (right). Were it not for the PS curvature sign reversal, almost no fusion would occur as the pore nucleation rate is much greater in the far bigger vesicle membrane.

In contrast, with 75% PE content all membranes have very high rupture thresholds as the fusion pore line tension τ is increased ∼2.5-fold due to the large negative curvature of PE that disfavors growth of positive curvature fusion pores (see *SI Text* and Fig. S2). Thus dead-end hemifusion is dominant (88%), with 12% fusion and almost no lysis due to the low vesicle tension. In the experiments no lysis was seen (10), in agreement with these predictions, but fusion and dead end occurred with equal probability, in contrast to the model predictions. We note that the model predicts equally probable fusion and dead end outcomes at a slightly lower PE content of ∼65% (Fig. 3B).

### Calcium-driven vesicle-planar membrane fusion stalls at hemifusion

During exocytosis, bioactive molecules are secreted by vesicle-plasma membrane fusion. Given the important role of calcium, an important question is whether non-specific *Ca*^2*+*^-driven fusion contributes to fusion in cells. Interestingly, *in vitro* studies showed that *Ca*^2*+*^fails to fuse giant vesicles with planar suspended bilayers, instead stalling at hemifusion (30, 31).

To address this unexplained finding, we applied our model to the vesicle-PM situation realized in the experiments of ref (30), using appropriate parameters (see Table 1). In agreement with these experiments, we find dead-end hemifusion is by far the most probable outcome (Fig. 4). The origin of this behavior is that the HD tension remains elevated for only ∼ 0.20 sec (eq. S32) due to its connection to the PM whose tension is relatively low. As a result, *<* 0.*1*% of vesicles fuse (eq. S35).

**Figure 4.**
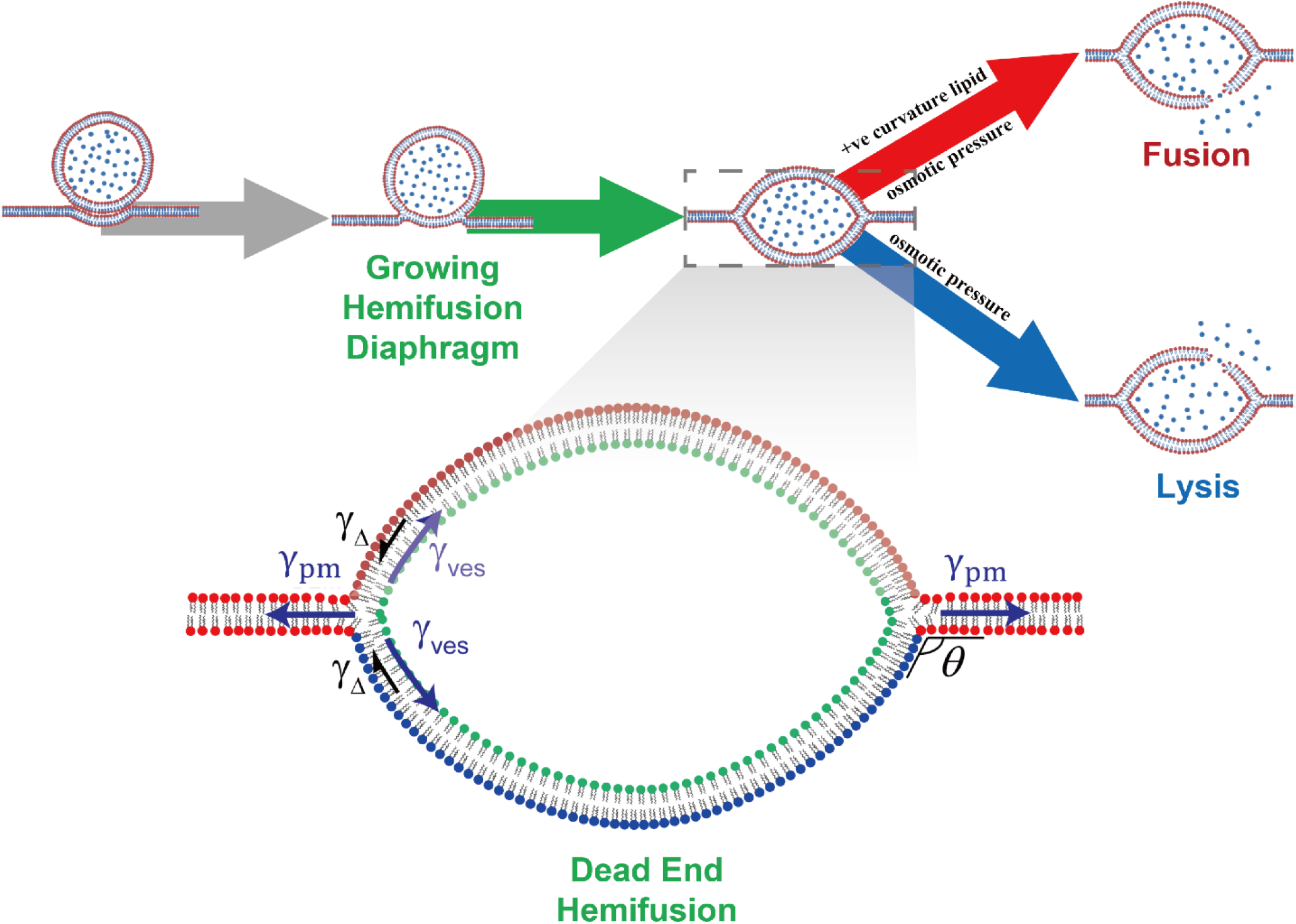
Calcium cannot typically fuse vesicles with a planar membrane (PM). Model predicted sequence is shown (schematic). At high *[Ca*^2*+*^], the vesicle has high tension and adheres strongly to the PM. Following hemifusion and HD growth the present model predicts that the vesicle tension is dissipated and a lens-shaped equilibrium hemifused complex is attained. Additional forces are required to drive fusion. Application of pore-promoting positive curvature lipids can selectively activate fusion or lysis.

The final equilibrium shape of the hemifused vesicle-PM complex is that of a symmetric lens with a contact angle *θ* (Fig. 4). In equilibrium, the vesicle must be sufficiently stretched in the plane of the PM that the force balance is satisfied, *γ*_pm_ = −2*γ*_ves_cos*θ*. Stretching of the vesicle is quite complex: the inner leaflet density decreases, while the outer leaflet compensates by drawing lipids from the PM, so its density increases (eqs. S39, S42). In consequence, the vesicle tension increases according to an effective stretch modulus *K*_eff_, depending on both the moduli for membrane tension *K* and for interleaflet tension *k*_Δ_ (see *SI Text*),

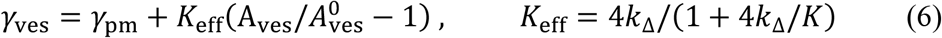

where *A*_ves_ is the vesicle area with initial value 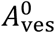. Now to leading order 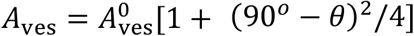, so the equilibrium lens area and angle are

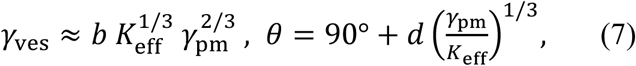

where *b* = 0.4, *d* = 72°. Using the parameters of Table 1, the equilibrium vesicle tension is 2.6 mN/m, well below the lysis threshold of 8.4 mN/m.

In ref. (30), LPC lysolipids were later added to the trans side of the PM, inducing fusion. Applying our model, their positive spontaneous curvature decreases the pore line tension and hence the tension threshold for lysis. We find the lysis threshold becomes as low as the equilibrium vesicle tension (2.6 mN/m) at a LPC mole fraction of ∼ 20%. Using the partitioning data of ref. (46), the 270 μM LPC solution used in refs. (30, 31, 47) is more than sufficient to achieve 20% LPC in the vesicle membrane, explaining why fusion was achieved.

## Discussion

### Calcium-mediated fusion is described by a network of pathways that pass through the hemifused intermediate

Here we showed that calcium-mediated fusion follows a network of pathways whose branch point is the hemifused intermediate (Fig 1B). The network is unchanging, but the conditions (Ca^2+^ concentration, lipid composition, membrane areas, geometry) regulate the frequency with which different pathways are selected (Figs 2A, 3B, 4). Pathway selection is made at the hemifused branch point and consists in selecting which membrane surface first ruptures by nucleation and dilation of a pore (43). Since the many-lipid pore dynamics are stochastic, the sampling of pathways in the network is stochastic, but the averages are fixed for a given set of conditions.

The fusion mechanism stems from powerful adhesion and contraction forces that divalent cations exert on charged membranes. These forces generate high membrane tensions and nucleate an expanding HD, a critical transition that channels two membrane tensions through the one HD bilayer (Fig. 1B, inset). However, even if the HD tension is high its rupture is not guaranteed because membranes can withstand super-rupture tensions for limited time periods (43). Since HD expansion progressively decreases tension, there is a small window in time when the HD may rupture (fusion); if the HD survives this transient, it grows to equilibrium (dead-end hemifusion, Figs. 2C, D and 3C, D). Another factor is membrane area, on which pore nucleation rates depend. The HD starts from zero and must grow before rupture is likely, while the vesicle area is much greater. Thus the vesicle may rupture first (lysis), despite its lower tension.

### Calcium concentration, lipid composition and membrane areas set the weightings of pathways in the network

We found that increasing Ca^2+^ concentration increases membrane tensions and rupture probability, so reducing the weighting for dead-end hemifusion (Fig. 2A, B). However the balance between fusion and lysis is more subtle since area comes into play: when all membranes are at super-rupture tensions, the larger vesicle membrane is more likely to rupture (lysis) as it nucleates more pores. Accordingly, fusion has maximum weighting at intermediate [Ca^2+^] and tension (Fig. 2A). Consistent with this, fusion was maximized at intermediate [Ca^2+^] in experiments using small vesicle bulk suspensions (6). Similarly, lipid composition radically influences the pathway weightings by altering the line tension of positive curvature membrane pores; for example, increased content of PE, a negative curvature lipid, increased the line tension, and the energy cost of pore expansion, with lowered fusion probability (Fig. 3).

### The effect of membrane geometry: calcium is ineffective at vesicle-planar membrane fusion

Ca^2+^-driven fusion assays using the vesicle-planar membrane geometry that arises during exocytosis are of particular interest. Surprisingly, calcium is an ineffective fusogen with this geometry (30, 31, 47), an unexplained fact. With typical experimental conditions our model reproduced the observed dead-end hemifusion: tensions dissipate too rapidly for fusion to occur, and the final state is an equilibrium lens-shaped low tension hemifused complex (Fig. 4).

Why is Ca^2+^ so effective at fusing vesicles (Figs. 2-3), yet cannot (without help from osmotic stress or lysolipids) fuse vesicles with a planar PM (Fig. 4)? There are two key factors. (i) PM tensions are not elevated by Ca^2+^-induced contraction, because the PM can draw lipids from the reservoir at its supporting torus to dissipate tension. Thus the initial HD tension is much lower than in vesicle-vesicle systems, where the HD supports two Ca^2+^-boosted vesicle tensions (Fig. 1, inset). (ii) The large PM is a virtually infinite lipid reservoir (48) unaffected by the hemifusion event. Thus, large Ca^2+^-induced vesicle tensions are dissipated on hemifusion with the PM, and we showed that the final equilibrium HD tension depends only on the relatively low tension of the PM (see eq. (**4**)).

### Cellular and calcium-mediated fusion may follow similar pathway networks

Fusion in cells may follow a network of fusion pathways centered on the hemifused intermediate similar to that for Ca^2+^-mediated fusion (Fig. 1B). Long-lived hemifused intermediates were detected on the fusion pathway in pneumatocytes and during yeast vacuole fusion (49, 50). Extended HDs have been observed between synaptic vesicles and the plasma membrane (∼5 nm in diameter), between granules and the plasma membrane of chromaffin cells (∼ 200 nm) and between yeast vacuoles (∼0.5-1 *μm*) (12, 13, 16, 51).

Thus, cellular fusion pathways may pass through hemifused intermediates with extended HDs. This is supported by data from reconstituted systems. In a study of fusion mediated by SNARE and synaptotagmin proteins, docked vesicle pairs either dead-end hemifused or remained docked (33). On introduction of Ca^2+^ some of the docked vesicles fused via a hemifused intermediate, fused directly or dead-end hemifused. Thus, higher fusion driving force (Ca^2+^ present) increased the fusion probability, similar to the pattern for Ca^2+^-driven fusion, Fig. 2A, B. Similarly, SNARE-mediated fusion rates were lower and dead-end hemifusion more probable for lower vesicle SNARE density (52) or when SNARE zippering was impeded by either hydrophobic molecules (53) or a mutation (54). In another study vesicles docked by SNAREs in point or extended contact adhered states evolved to dead-end hemifusion or fusion (32). The presence of PE in vesicle membranes reduced the incidence of SNARE-mediated fusion of vesicles with supported bilayers, produced significant dead-end hemifusion and decreased the hemifusion to fusion transition rate (55), similar to our predictions for the pore growth rate *k*_grow_ (eq. (**5)** and Fig. 3). Finally, hemifusion is a productive intermediate on the pathway to fusion mediated by influenza’ s fusion protein hemagglutinin, and can also be a dead-end state (56-58).

These parallels suggest that the network of fusion pathways may be intrinsic to the lipid bilayers themselves, independently of the fusogen involved. Different fusogens may generate different outcome distributions and pathway kinetics due to their differing force-generating mechanisms. Although SNAREs, hemagglutinin and calcium generate force via different mechanisms, they must all contend with the same forces that resist progress along the pathway to membrane fusion, set by the biophysical properties of the phospholipid bilayers.

## Author contributions

B.O’ S. and J. W. designed the research; J.W, B.O’ S., D.A. and B.S.S. performed the research and analyzed the data; B.O’ S, J.W and D.A. wrote the paper.

## Acknowledgements

Research reported in this publication was supported by the National Institute of General Medical Sciences of the National Institutes of Health under award number R01GM117046 to B.O. The content is solely the responsibility of the authors and does not necessarily represent the official views of the National Institutes of Health. We thank Andreas Hermann and Joerg Nikolaus for helpful discussions and access to their data prior to submission.

