## Supporting Information for "A hemifused complex is the hub in a network of pathways to membrane fusion"

b: St. Luke's School, New Canaan, CT 06840

February 15, 2022

### Time evolution of membrane tensions

The fusion rate constants featuring in eq. (5) of the main text depend on the time-dependent HD area and membrane tensions. We previously produced a model to calculate these. In this section, we give brief descriptions of HD growth mechanisms and the corresponding calculations. For more details, please see refs (1, 2).

**Changes in membrane tension due to vesicle-vesicle adhesion and HD growth.** It is well known that the membrane tension of the round portion of the vesicle decreases linearly with a decrease of the total membrane area,

$$\gamma_{\mathrm{ves}}(t) = \gamma^0 + K\left(\frac{A_{\mathrm{tot}}(t)}{A_{\mathrm{tot}}^0} - 1\right) , \qquad \text{S1}$$

where *K*=265 mN/m is the bilayer stretch modulus, $\gamma^0$ is the vesicle tension before adhesion (will discuss this parameter later) and $A_{\mathrm{tot}}^0$ is the total area of the pair of vesicles before adhesion

$$A_{\mathrm{tot}}^0 = 2 \times (4\pi R_{\mathrm{ves}}^2) \qquad \text{S2}$$

Where $R_{\mathrm{ves}}$ is the unadhered vesicle radius. It can be shown using simple spherical geometry that the surface area of the round portion of each vesicle is

$$A_{\mathrm{ves}}(t) = 2\pi R(t)^2(1 + \cos\theta(t)) \qquad \text{S3}$$

Where $\theta(t)$ is the the vesicle-vesicle contact angle during fusion, $R(t)$ is the radius of the spherical portion of the vesicle during fusion, and the adhered flat region has an area

$$A_{\mathrm{ad}}(t) = \pi R(t)^2 \sin^2\theta(t) - A_{\mathrm{HD}}(t), \qquad \text{S4}$$

where $A_{\mathrm{HD}}(t)$ is the HD area (Fig. S1). Thus, the total area for an adhered vesicle pair is that

$$A_{\mathrm{tot}}(t) = 2\big(A_{\mathrm{ves}}(t) + A_{\mathrm{ad}}(t)\big) + A_{\mathrm{HD}}(t) \qquad \text{S5}$$

Substituting eqs. S2-S5 into eq. S1, we have the vesicle membrane tension $\gamma$ as a function of the contact angle, vesicle radius, and HD area,

$$\gamma_{\mathrm{ves}} = \gamma^0 + K\left(\frac{4\pi R^2(1+\cos\theta) + 2\pi R^2\sin^2\theta - A_{\mathrm{HD}}}{2A_0} - 1\right) \qquad \text{S6}$$

Now, as the contact angle $\theta(t)$ increases during vesicle flattening due to adhesion, the vesicle radius must grow to conserve vesicle volume according to the geometric relationship between vesicle radius, volume, and contact angle,

$$V_{\text{ves}}(t) = \frac{\pi}{3} R(t)^3 (2 + 3\cos\theta(t) - \cos^3\theta(t)) \qquad \text{S7}$$

Finally, the relationship between contact angle and membrane tension is due to the force balance (Young's equation) along the perimeter of the contact zone (Fig. S1)

$$\cos\theta(t) = 1 - \frac{W}{2\gamma_{\text{ves}}(t)} \quad (\text{vesicle} - \text{vesicle adhesion}), \qquad \text{S8}$$

where $W$ is the vesicle-vesicle adhesion energy. We will discuss it in the later section. Now, given the vesicle volume, for each value of $A_{\text{HD}}(t)$ eqs. S6-S8 can be solved simultaneously to yield $\theta$, $R$, and $\gamma$.

The above describes the procedure for vesicles in suspension, such as in the experiments of Rand et al. and Kachar et al. The procedure is the same for the experiments of Nikolaus et al., except the force balance (eq. S8) is slightly changed since the tension of one, not two, vesicle attempts to separate the membrane from the zone of adhesion with the glass coverslip substrate,

$$\cos\theta(t) = 1 - \frac{W}{\gamma_{\text{ves}}(t)} \quad (\text{vesicle} - \text{substrate adhesion}). \qquad \text{S9}$$

**Estimation of the vesicle-vesicle adhesion energy $W$ and vesicle tension $\boldsymbol{\gamma^0}$ in the adhered state for the experiments of refs. (3) and (4).** To estimate the vesicle-vesicle adhesion energy for DOPS vesicles, we first measured the vesicle-vesicle contact angles in the microscopy images of .ures 2B and 2E in Reese and Rand (3) to be on average $\theta \approx 40°$. We assumed the images to correspond to the adhered state (see Fig. 1A of main text) or an early hemifused state with a small HD, and that negligible vesicle leakage had occurred between the addition of $Ca^{2+}$ and image capture. We then simultaneously solved eqs. S6-S8 using $\theta = 40°$, $A_{\text{HD}} = 0$, and $\gamma^0 = 9.4$ mN/m for the adhesion energy W=5.3 mN/m and the adhered vesicle tension of the rounded region is $\gamma_{\text{ves}} = 11.1$ mN/m.

For the DOPS/DOPE mixtures of Kachar et al., we measured similar contact angles $\theta \approx 40°$ in their figures 2B and 3A. (We excluded the image in their Fig. 1A which shows a smaller vesicle strongly deformed in adhesion with a larger but less deformed vesicle adhered with significantly larger contact angles suggesting that the smaller vesicle had sufficient time to leak.) Since the $Ca^{2+}$-induced membrane tension varies only slightly between ~8.0 and 9.4 mN/m, that the contact angles were similar for various compositions suggests adhesion energy is somewhat weakly dependent on DOPS/DOPE composition ratios. Hence for simplicity, for all DOPS/DOPE compositions, we assumed that the adhesion energy is effectively independent of composition, W=5.3 mN/m.

**Reduction of membrane tension due to leakage.** Since the pressure inside the vesicle is an amount $\Delta P$ higher than that outside the vesicle, water leaks through the membrane at a volumetric rate

$$\dot{V}_{\rm ves}(t) = \frac{\eta v_{\rm w}}{kT} A_{\rm ves} \Delta P(t) \qquad \text{S10}$$

where η is the membrane permeability and $v_{\rm w} = 3{\rm x}10^{-11}$ μm$^3$ is the molecular volume of water. We take $\eta = 12.5$ μm/s (5). Inserting eq. S10 into the Young-Laplace relation between tension and pressure, $\Delta P = 2\gamma_{\rm ves}/R$ and approximating $A_{\rm ves} \approx 4\pi R^2$ ,

$$\dot{V}_{\rm ves} = \frac{8\pi\eta v_{\rm w}}{kT} R\gamma \qquad \text{S11}$$

In the calculations, vesicle volume was updated after each time step of duration Δt, reduced by amount $\dot{V}\Delta t$. Accordingly, in calculating the tension for each time step, eq. S6 was updated with the new, reduced vesicle volume.

#### Kinetic model of calcium-mediated membrane fusion

The branched pathway schematized in eq. (5) and Fig. 1B of the main text corresponds formally to a set of coupled differential equations which describe the branched pathway to fusion, lysis or dead-end hemifusion. Here we derive these equations in a form which can be solved numerically. Let us first consider the path to fusion via HD rupture. Consider an ensemble population of *N* interacting vesicle pairs. At time zero the population of the adhered state **A** is *A*(0)=*N*. As time goes on the populations *A*, *HD*, *HD** and *F* of the intermediate and outcome states **A**, **HD**, **HD*** and **F**, respectively, evolve according to

$$\begin{aligned}
\frac{dA}{dt} &= -k_{\text{hemi}}\, A \,-\, \frac{A}{A+HD+HD^*}\, k_{\text{grow}}^{\text{ves}}(t)V^* \\
\frac{dHD}{dt} &= k_{\text{hemi}}A - k_{\text{nuc}}^{\text{HD}}(t)HD + k_{\text{seal}}HD^* \,-\, \frac{HD}{A+HD+HD^*} k_{\text{grow}}^{\text{ves}}(t)V^* \\
\frac{dHD^*}{dt} &= k_{\text{nuc}}^{\text{HD}}(t)HD - \left(k_{\text{seal}} + k_{\text{grow}}^{\text{HD}}\right)HD^* \,-\, \frac{HD^*}{A+HD+HD^*} k_{\text{grow}}^{\text{ves}}(t)V^* \\
\frac{dF}{dt} &= k_{\text{grow}}^{\text{HD}}HD^* \; .
\end{aligned} \qquad \text{S12,}$$

where $k_{\text{hemi}}$ is the HD nucleation rate.

Above, all terms are apparent from eq. 5 of the main text except those proportional to *V** which account for destruction of **HD** and **HD*** intermediates due to the vesicle lysis "side reaction." Similarly, for vesicle surfaces, the state populations evolve according to

$$\begin{aligned}
\frac{dV}{dt} &= -k_{\text{nuc}}^{\text{ves}}(t)V + k_{\text{seal}}V^* - \frac{V}{V+V^*} k_{\text{grow}}^{\text{HD}}(t)HD^* \\
\frac{dV^*}{dt} &= k_{\text{nuc}}^{\text{ves}}(t)V - \left(k_{\text{seal}} + k_{\text{grow}}^{\text{ves}}(t)\right)V^* - \frac{V^*}{V+V^*} k_{\text{grow}}^{\text{HD}}(t)HD^* \\
\frac{dL}{dt} &= k_{\text{grow}}^{\text{ves}}(t)V^*.
\end{aligned} \qquad \text{S13}$$

Again, all terms are apparent from eq. (5) of the main text except those proportional to *HD** which account for destruction of vesicle pairs in the **V** and **V*** states whose HD ruptures before lysis occurs.

We solved eqs. S12-S13 for the relative probabilities of these three outcomes as a function of calcium concentration, vesicle size, and lipid composition. The rate constants for each step in eqs. S1-S2 are described in the main text. They are developed from Evans et al. (6) and depend on the evolving areas and membrane tensions of the expanding HD and vesicle membranes.

#### Estimation of Parameter Values in Table 1 of the Main Text

**Estimation of divalent cation-induced lipid condensation.** For the lipid mixtures studied in this paper, we estimate the total cation shrinkage factor $\epsilon_{\text{cat}}$ based on those for the pure lipid species PC, PS, and PE. We name those corresponding single-component shrinkage factors $\epsilon_{\text{PC}}$,

$\epsilon_{\mathrm{PS}}$ and $\epsilon_{\mathrm{PE}}$, respectively, and estimate these values from previous measurements. Assuming linear dependence on the composition fractions $\chi_i$ for each species $i$, we have

$$\epsilon_{\mathrm{cat}} = \chi_{\mathrm{PS}}\epsilon_{\mathrm{PS}} + \chi_{\mathrm{PC}}\epsilon_{\mathrm{PC}} + \chi_{\mathrm{PE}}\epsilon_{\mathrm{PE}}. \qquad \text{S14}$$

*PS condensation.* For anionic PS lipids, we use the measurements of divalent-induced tensions $\gamma_{\mathrm{mono}}$ for PS monolayers of Ohki et al. (7). At each [$Ca^{2+}$] value and [$Mg^{2+}$] value where monolayer tension was measured, we estimated $\epsilon_{\mathrm{PS}} = \gamma_{\mathrm{mono}}/K_{\mathrm{mono}}$ where we take the monolayer stretch modulus $K_{\mathrm{mono}}$ as half the bilayer stretch modulus, $K_{\mathrm{mono}} = K/2$. The same procedure was used in refs. (1) and (2). We take $K$=265 mN/m (8).

*PC condensation.* For PC lipids, quantitative data within the literature for Ca-induced condensation is inconsistent. Huster et al found 1.4% areal condensation of DOPC in 5 mM Ca2+ (relative to Ca2+-free solutions) (9). Uhrikova et al. found much stronger areal condensation, 7.1% for DPPC in 5 mM $Ca^{2+}$ (10). Higher values of [$Ca^{2+}$] were found by Uhrikova et al. to reduce condensation with increasing Ca2+. The authors attributed this surprising effect to large molecular rearrangements of the lipids, perhaps due to a lipid phase change. Since phase changes are not observed in DOPC/DOPS/DOPE mixtures in Ca2+ (9), we ignored the Uhrikova data for concentrations greater than 10 mM. Given the inconsistent literature data, we chose to fit the data of Uhrikova et al. (10) between 1 and 10 mM [$Ca^{2+}$] to a logarithm, $\epsilon_{\mathrm{PC}} = 0.013 \ln[\mathrm{Ca}^{2+}] + 0.038$. Our choice of function was driven by assuming that PC behaves qualitatively similarly to PS, for which in the experiments of Ohki et al. the tension increased roughly logarithmically for concentrations $\lesssim$ 10 mM. Lacking Mg-specific data for PC, we assumed that it responds identically to Mg and Ca.

*PE condensation.* PE does not form vesicles at physiological conditions, and thus pure PE condensation data does not, to the best of our knowledge, exist. Since PE and PC headgroups are both zwitterionic, we assume that $\epsilon_{\mathrm{PC}} = \epsilon_{\mathrm{PE}}$. This is the same procedure used in refs. (1) & (2).

**Bilayer tension before hemifusion.** For the lipid mixtures considered in this work we estimated the initial membrane tensions induced by calcium, $\gamma^0$, using the monolayer tension (only outer leaflets are bathed in cation solutions in fusion experiments) for each pure lipid species under similar conditions, and assumed linear composition dependence:

$$\gamma^0 = K_{\mathrm{mono}}\epsilon_{\mathrm{cat}} \qquad \text{S15}$$

By combing S14 and S15, we get

$$\gamma^0 = \chi_{\mathrm{PS}}\gamma_{\mathrm{PS}} + \chi_{\mathrm{PC}}\gamma_{\mathrm{PC}} + \chi_{\mathrm{PE}}\gamma_{\mathrm{PE}}. \qquad \text{S16}$$

For PS lipids we took $\gamma_{\mathrm{PS}}$ directly from the Ohki et al data for each [$Ca^{2+}$]. For PC lipids we took $\gamma_{\mathrm{PC}} = K_{\mathrm{mono}}\,\epsilon_{\mathrm{PC}}$, and again lacking PE-specific data we took $\gamma_{\mathrm{PE}} = \gamma_{\mathrm{PC}}$.

**Critical tension of nascent pores.** The rates of fusion by HD rupture and vesicle lysis (ie $k_{\mathrm{grow}}^{\mathrm{HD}}(t)$ and $k_{\mathrm{grow}}^{\mathrm{ves}}(t)$ in eq. (5) of the main text) depend on both the respective membrane tensions $\gamma(t)$ and the critical tension $\gamma_{\mathrm{crit}}$. Expansion of nascent pore lowers the total energy by decreasing the total area, though only for large pores above a critical radius (6, 11). The total energy of a membrane with a nascent pore is

$$E(r_{\mathrm{pore}}) = 2\pi r_{\mathrm{pore}}\tau - \pi r_{\mathrm{pore}}^2\gamma, \qquad \text{S17}$$

where $r_{\mathrm{pore}}$ is the pore radius (Fig. S2). The energy maximum occurs at a critical pore radius $r_{\mathrm{c}} = \tau/\gamma$. After a pore grows to this critical size through thermal fluctuations it will grow spontaneously. Equating eq. S16 to the thermal energy $kT$, the critical tension is

$$\gamma_{\mathrm{crit}} = \pi\tau^2/kT. \qquad \text{S18}$$

Thus, to calculate the critical tension for a given membrane we must estimate the line tension (see directly below).

The rupture theshold tension. The rupture threshold $\gamma_{\mathrm{rupture}}$, where pore growth and resealing rates are equal. So

$$k_{\mathrm{grow}} = k_{\mathrm{grow}}^{\mathrm{o}}\left(\gamma_{\mathrm{rupture}}/\gamma_{\mathrm{crit}}\right)^{1/2} \mathrm{e}^{-\gamma_{\mathrm{crit}}/\gamma_{\mathrm{rupture}}} = k_{\mathrm{seal}}. \qquad \text{S19}$$

**Calculation of pore line tension $\boldsymbol{\tau}$ and the effect of lipid partitioning**. The pore line tension is strongly dependent on the spontaneous curvature of the component lipids. Here in our calculation of the line tension we consider the partitioning of positive (negative) spontaneous curvature lipids into (away from) the positively curved pore. This lowers the line tension considerably as compared to a simpler model where the pore composition matches the "bulk" membrane composition.

Consider a nascent formed in a bilayer of thickness $2\delta$ (Fig. S2). For lipid species *i*, the bending energies per lipid in the pore and bulk (non-pore portion of membrane) are, respectively,

$$\epsilon_{\mathrm{i}}^{\mathrm{P}} = \frac{\kappa}{2}\rho_{\mathrm{lip}}^{-1}(C^{\mathrm{P}} - C_{\mathrm{i}}^{\mathrm{o}})^2$$
$$\epsilon_{\mathrm{i}}^{\mathrm{B}} = \frac{\kappa}{2}\rho_{\mathrm{lip}}^{-1}(C^{\mathrm{B}} - C_{\mathrm{i}}^{\mathrm{o}})^2 \qquad \text{S20}$$

where $\kappa$ is the monolayer bending modulus, $\rho_{\mathrm{lip}}$ is the areal density of lipids in the bilayer, $C^{P}$ and $C^{B}$ are the curvatures of the pore and planar bilayer respectively, and $C_{\mathrm{i}}^{\mathrm{o}}$is the spontaneous curvature of species *i*. Note eq. S18 above follows the standard Helfrich model where bending energy is proportional to the square of the curvature strain (12). We estimate that the pore curvature is the inverse of the monolayer thickness, $1/\delta$, and the bulk vesicle and HD bilayers have zero curvature. Spontaneous curvatures of various lipid species are shown in Table S1. We take $\kappa$=10 *kT* and $\rho_{\mathrm{lip}}$=1.67 $nm^{-2}$. Thus, the line tension of pure lipid species *i* is

$$\tau_{\mathrm{i}} = \rho_{\mathrm{lip}} l_{\mathrm{pore}}\left(\epsilon_{\mathrm{i}}^{\mathrm{P}} - \epsilon_{\mathrm{i}}^{\mathrm{B}}\right) \qquad \text{S21}$$

where $l_{\mathrm{pore}}$ is the cross-sectional half-circumference of the pore, as schematized in Fig. S2. We ensured that this model predicts the correct line tension of 11.5 pN measured previously (6) for pure PC bilayers. Thus we calibrated eq. S20 above by setting $\tau = 11.5$ pN and solving for $l_{\mathrm{pore}} = 1.3$ nm. We retain this calibrated value for subsequent calculations for 2-component membranes.

In multi-component membranes, lipid species will favor residing in the pore (bulk membrane) region if it has a positive (negative) spontaneous curvature. For a composition of 2 lipid types, labeled 1 and 2, the total free energy of the pore and bilayer is

$$G = N_{\mathrm{tot}}^{\mathrm{P}}(\phi_1^{\mathrm{P}}\mu_1^{\mathrm{P}} + \phi_2^{\mathrm{P}}\mu_2^{\mathrm{P}}) + N_{\mathrm{tot}}^{\mathrm{B}}(\phi_1^{\mathrm{B}}\mu_1^{\mathrm{B}} + \phi_2^{\mathrm{B}}\mu_2^{\mathrm{B}}) \qquad \text{S22}$$

where $\phi_{\mathrm{i}}$ is the mole fraction of species *i* in either the pore (P) or the bulk (B), and $\mu_{\mathrm{i}}$ is the free energy per lipid of species *i* in the pore or the bulk. $N_{\mathrm{tot}}$ is the total number of lipids in each region. For a given pore radius, assuming incompressibility, the total number of lipids in the pore is fixed, but the species may exchange with the bulk. The realized composition is that which minimizes the total free energy of eq. S22. Taking the derivative of *G* above, this occurs when

$$\Delta\mu_1 = \Delta\mu_2\,, \quad \Delta\mu_\mathrm{i} \equiv \mu_\mathrm{i}^\mathrm{P} - \mu_\mathrm{i}^\mathrm{B}. \qquad \text{S23}$$

The above result allows us to write the line tension of the mixture in terms of the chemical potential of species 1,

$$\tau = \rho_\mathrm{lip} l_\mathrm{pore} \frac{dG}{dN_\mathrm{tot}^\mathrm{P}} = \rho_\mathrm{lip} l_\mathrm{pore} \Delta\mu_1. \qquad \text{S24}$$

Now to calculate the lipid free energy we use the bending energy of eq. S20 and assume ideal gas entropy,

$$\begin{aligned} \Delta\mu_1 &= \Delta\epsilon_1 + kT\ln(\phi_1^\mathrm{P}/\phi_1^\mathrm{B}) \\ \Delta\mu_2 &= \Delta\epsilon_2 + kT\ln(\phi_2^\mathrm{P}/\phi_2^\mathrm{B}) \end{aligned} \qquad \text{S25}$$

Thus using $\Delta\mu_1 = \Delta\mu_2$ (eq. S23) we solve for the pore composition $\phi_1^\mathrm{P}$ as a function of the bulk composition $\phi_1^\mathrm{B}$,

$$\frac{\phi_1^\mathrm{P}}{1-\phi_1^\mathrm{P}} = \frac{\phi_1^\mathrm{B}}{1-\phi_1^\mathrm{B}} e^{-\epsilon/kT}, \quad \epsilon \equiv \Delta\epsilon_1 - \Delta\epsilon_2 \qquad \text{S26}$$

where $\epsilon$ is the relative energetic advantage of species 2 being in the pore vs. species 1. Note that $\epsilon > 0$ if species 2 is porephilic and species 1 is porephobic. Thus, we insert eq. S26 into eq. S25, and that into eq. S22, we obtain

$$\frac{\tau}{\rho_\mathrm{lip} l_\mathrm{pore}} = \Delta\epsilon_1 - kT\ln\left\{\phi_1^\mathrm{B} + (1-\phi_1^\mathrm{B})e^{-\epsilon/kT}\right\}. \qquad \text{S27}$$

Hence we use eq. S27 to calculate the line tension of 2-component membranes as a function of their bulk composition fractions and spontaneous curvatures.

Eq. S27 is valid for any 2-component membrane. This result could be extended to a much more complicated general result for any multi-component mixture of *n* species. However, for simplicity, when considering the 3-component mixture of Nikolaus et al we used the line tension procedure above with a minor modification. We took species 1 to be PE since its curvature is the most extreme. For the properties of lipid species 2, we used the composition –weighted average of the PC and PS properties since these lipids have relatively mild spontaneous curvatures.

Similarly for LPC in asolectin (GUV-PM experiments) we took LPC to be lipid species 1 and the asolectin mixture to be species 2.

#### GUV-PM fusion

**Initial HD growth kinetics (GUV-PM fusion).** We showed previously (2) that HD areal growth is initially linear in time for times much smaller than the characteristic "diffusion" time for relaxation of interleaflet tension over the vesicle, $\sim R_{\text{ves}}^2 \lambda / k_\Delta$. Modifying the symmetric vesicle-vesicle situation of our previous work to the asymmetric system where the vesicle and PM have differing tensions, we have

$$A_{\text{HD}} = \alpha t, \qquad \alpha = c \frac{f_{\text{therm}}}{\ln 1/f_{\text{therm}}} \frac{k_\Delta}{\lambda}, \qquad f_{\text{therm}} = \frac{\gamma_{\text{PM}} + \gamma_0}{4k_\Delta} + \frac{\epsilon_{\text{cation}}}{2}. \qquad \text{S28}$$

Here $c$=10.6 is a numerical constant (2) and $\alpha$ is the initial HD areal growth rate.

**Relaxation of HD tension during HD growth (GUV-PM fusion).**

For small HDs ($A_{\text{HD}} \ll A_{\text{ves}}$, the HD tension resulting from a force balance is

$$\gamma_{\text{HD}} = \gamma_{\text{PM}} + \gamma_{\text{ves}}, \qquad \text{S29}$$

where $\gamma_{\text{PM}}$ is fixed but $\gamma_{\text{ves}}$ decays with HD growth according to

$$\gamma_{\text{ves}} = \gamma_0 - K \frac{A_{\text{HD}}}{2A_{\text{ves}}^{\text{o}}}, \qquad \text{S30}$$

which results from applying eq. S for the case of negligibly weak adhesion and negligibly slow water leakage. Hence combining eqs. S28 and S30, the HD tension decays linearly in time,

$$\gamma_{\text{HD}} = \gamma_{\text{PM}} + \gamma_0 - \frac{K\alpha}{2A_{\text{ves}}^{\text{o}}} t \ . \qquad \text{S31}$$

It follows that the HD tension relaxes to safe sub-lysis levels below the rupture threshold $\gamma_{\text{rupture}}$ by the time

$$\tau_{\text{relax}} = \frac{2A_{\text{ves}}^{\text{o}} (\gamma_{\text{PM}} + \gamma_0 - \gamma_{\text{rupture}})}{K\alpha} \qquad \text{S32}$$

**Initial fusion kinetics (GUV-PM fusion).**

Let us calculate an upper bound to the amount of fusion that occurs in GUV-PM systems. To do this, we assume that HD tension is sufficiently high that once a pore is nucleated, it immediately grows without resealing. Hence nucleation rate-limited fusion rate is

$$\frac{dF}{dt} = k_{\text{nuc}}^{\text{HD}}. \qquad \text{S33}$$

Now combining this with $k_{\text{nuc}}^{\text{HD}} = \text{A}_{\text{hd}}\, r_{\text{nuc}} \text{e}^{\gamma_{\text{hd}}/\gamma^*}$ and eq. S33 of SI and using the HD tension of eq. S31, we have the time-dependent fusion rate

$$\frac{dF}{dt} = \alpha r_{\text{nuc}} e^{(\gamma_0+\gamma_{\text{PM}})/\gamma^*} t. \qquad \text{S34}$$

Hence the fusion rate is initially zero since the nascent HD has zero area, but the rate grows linearly in time as does the HD area. To determine upper bound for the amount of fusion that occurs before tension relaxes to safe levels, we integrate eq. S34 from time zero to t=$\tau_{\text{relax}}$,

$$F_{\text{max}} = \frac{1}{2} \alpha r_{\text{nuc}} e^{(\gamma_0+\gamma_{\text{PM}})/\gamma^*} \tau_{\text{relax}}^2. \qquad \text{S35}$$

**Hemifusion equilibrium (GUV-PM).**

Here we derive the equilibrium results for hemifusion of a vesicle with a PM presented in eq. (**7**) of the main text. Now PM tensions are normally much less than the interleaflet modulus $k_\Delta$ or the stretch modulus $K$. We will see this leads to equilibrium where the vesicle has a lens shape only slightly distorted from a sphere. Mathematically, $\gamma_{\text{pm}}/k_\Delta$ and $\gamma_{\text{PM}}/K$ are small. We first must identify what criteria determine hemifusion equilibrium. Now as argued in the main text the hemifused state adopts a symmetric lens configuration (see Fig. **4**) where the non-HD and HD regions are spherical caps as in the configuration of a liquid drop at an interface (13).

By symmetry of the lens structure, we get

$$\gamma_{\text{HD}} = \gamma_{\text{ves}}. \qquad \text{S36}$$

According to the general relation of the tension for three-branch membrane structure in ref (1), the relation between the tension of the planar membrane, that of the vesicle membrane tension, and the interleaflet tension is that

$$\gamma_{\mathrm{pm}} \approx \gamma_{\mathrm{ves}} + 2\gamma_{\Delta}. \qquad \text{S37}$$

For the derivation of eq. S37, please see ref. (1). Substituting $\gamma_{\mathrm{ves}} \approx K\left(1-\frac{\bar{\rho}}{\rho_0}\right)$, $\gamma_{\mathrm{pm}} \approx K\left(1-\frac{\rho_{\mathrm{pm}}}{\rho_0}\right)$, and $\gamma_{\Delta} \approx k_{\Delta}\frac{\rho_{\mathrm{in}}-\rho_{\mathrm{out}}}{\rho_0}$ in eq. S37, where $K$ and $k_{\Delta}$ are bilayer stretch modulus and interleaflet modulus respectively ($\bar{\rho}$ is the mean lipid density of the vesicle membrane), we get

$$K\left(\frac{\bar{\rho}-\rho_{\mathrm{pm}}}{\rho_0}\right) \approx 2k_{\Delta}\frac{\rho_{\mathrm{in}}-\rho_{\mathrm{out}}}{\rho_0}. \qquad \text{S38}$$

Substituting $\bar{\rho} \equiv \frac{\rho_{\mathrm{in}}+\rho_{\mathrm{out}}}{2}$ into eq. S38, we get

$$\frac{\rho_{\mathrm{in}}-\rho_{\mathrm{out}}}{\rho_{\mathrm{in}}-\rho_{\mathrm{pm}}} \approx \frac{K_{\mathrm{eff}}}{2k_{\Delta}}, \quad K_{\mathrm{eff}} = \frac{Kk_{\Delta}}{k_{\Delta}+K/4}. \qquad \text{S39}$$

By substituting eq. S39 into the interleaflet tension $\gamma_{\Delta} \approx k_{\Delta}\frac{\rho_{\mathrm{in}}-\rho_{\mathrm{out}}}{\rho_0}$, we get

$$\gamma_{\Delta} \approx \frac{K_{\mathrm{eff}}}{2}\left(\frac{\rho_{\mathrm{in}}-\rho_{\mathrm{pm}}}{\rho_0}\right). \qquad \text{S40}$$

Substituting eq. S40 into eq. S39, we get

$$\gamma_{\mathrm{pm}} - \gamma_{\mathrm{ves}} \approx K_{\mathrm{eff}}\left(\frac{\rho_{\mathrm{in}}-\rho_{\mathrm{pm}}}{\rho_0}\right). \qquad \text{S41}$$

where $K_{\mathrm{eff}}$ is an effective stretch modulus of the vesicle membrane with the outer leaflet connected with a large membrane reservoir.

For the inner leaflet of the vesicle, by assuming volume conservation and total lipid number conservation, we get the relative change of the lipid density of the inner leaflet of the vesicle is that

$$\frac{\rho_{\mathrm{in}}}{\rho_0} \approx 1-\frac{\phi^2}{4} \qquad \text{S42}$$

based on vesicle volume conservation and inner leaflet lipid number conservation. Substituting eq. S42 and $\frac{\rho_{\mathrm{pm}}}{\rho_0} \approx 1-\frac{\gamma_{\mathrm{pm}}}{K}$ into eq. S41, we get

$$\gamma_{\mathrm{ves}} \approx \gamma_{\mathrm{pm}}\left(1-\frac{K_{\mathrm{eff}}}{K}\right) + K_{\mathrm{eff}}\frac{\phi^2}{4}. \qquad \text{S43}$$

According to force balance for the tension of vesicle membrane and planar membrane base on the geometry of the lens junction a lens, we get

$$\gamma_{\text{pm}} = 2\gamma_{\text{ves}} \sin\phi \approx 2\gamma_{\text{ves}}\phi. \qquad \text{S44}$$

Substituting $\gamma_{\text{ves}} \approx \frac{\gamma_{\text{pm}}}{2\phi}$ into eq. S43, we get

$$\phi^3 + \frac{4\gamma_{\text{pm}}\left(1 - \frac{K_{\text{eff}}}{K}\right)}{K_{\text{eff}}}\phi - \frac{2\gamma_{\text{pm}}}{K_{\text{eff}}} \approx 0. \qquad \text{S45}$$

By solving eq. S45, the approximation solution to the leading order is

$$\phi \approx \left(\frac{2\gamma_{\text{pm}}}{K_{\text{eff}}}\right)^{1/3}. \qquad \text{S46}$$

Substituting eq. S46 into eq. S44, we get

$$\gamma_{\text{ves}} \approx \left(\frac{K_{\text{eff}}}{16}\right)^{1/3} \gamma_{\text{pm}}^{2/3}. \qquad \text{S47}$$

This indicates that the equilibrium tension of the vesicle is governed by both the planar membrane tension and the effective stretch modulus $K_{\text{eff}}$ of the membrane.

**Calculation of osmotic pressure in GUV-PM fusion.** We calculated the osmotic pressure used in refs. (14) & (15) by converting the osmolarity difference between the vesicle contents and the solution reported in each study to an osmotic pressure. Then, using Young's equation, $\gamma = P_{\text{osm}}R_{\text{ves}}/2$, we calculate the new membrane tension. To determine if fusion or lysis occur or if instead dead-end hemifusion is stable, we compared this tension to the rupture tension (determined by setting $k_{\text{grow}}^{\text{HD}} = k_{\text{seal}}$ as described in the main text). Because the ves-PM hemifused state is symmetrical (Fig. 4), fusion and lysis are equally likely unless osmotic conditions are adjusted to be asymmetric across the PM to favor fusion over lysis. We take the vesicle radius to be the same as the radius prior to hemifusion, consistent with the small changes to vesicle radius and area upon hemifusion we find above (see SI section *Hemifusion equilibrium (GUV-PM)*).

**Calculation of effects of lysolipids in GUV-PM fusion.**

In the experiments of Chernomordik et al (14), a solution of 270 μM lysolipid (LPC) was added to the opposite ('trans') side of the PM from that where the GUVs resided ('cis' side). LPC then partitions itself between the soluble and membrane phases (the LPC composition of the membrane increases with the soluble LPC concentration). LPC has strongly positive spontaneous curvature and thus reduces the pore line tension and the threshold membrane tension for rupture. Our approach was to calculate the LPC solution concentration required to drive fusion by reducing the membrane rupture tension below the HD membrane tension (we showed above that $\gamma_{\mathrm{HD}}$=2.6 mN/m for GUVs dead-end-hemifused with the PM).

First we estimated the spontaneous curvature of the GUVs and PMs of ref. (14) composed of a soy bean extract lipid mixture, called asolectin, was -0.13 nm$^{-1}$. Note this lies between that for DOPC (-0.11 nm$^{-1}$) and for DOPE (-0.35 nm$^{-1}$), which are the major constituents of asolectin (Asolectin is a natural soybean mixture which contains ~31% PC, ~28% PE, ~22% PI and PA combined, ~12% neutral lipids, ~3% lysolipids, and ~5% "other lipids" (14)). The spontaneous curvature of LPC is +0.26 nm$^{-1}$ (16). Hence we used the procedure of the SI section *Calculation of pore line tension τ and the effect of lipid partitioning* to calculate the pore line tension for various ratios of LPC to asolectin. From the line tensions for each composition we could then determine the rupture tension by setting the pore growth and sealing rates equal, $k_{\mathrm{grow}}^{\mathrm{HD}} = k_{\mathrm{seal}}$. We found that ~20% LPC is required to reduce the rupture tension to 2.6 mN/m such that the HD tension could rapidly drive complete fusion.

To determine the soluble LPC concentration required to produce 20% LPC in the membrane, we relied on the partitioning data of Needham and Zhelev (17). They found that soluble LPC concentrations of 1 μM, 5 μM, and 10 μM led to membrane compositions that were 2%, 7%, and 15% LPC, respectively. Extrapolating this data suggests that ~13 μM LPC will produce sufficient LPC in the membrane to drive fusion. Hence our model predictions are consistent with the experimental observation: the 270 μM LPC used by Chernomordik et al is more than sufficient to drive fusion (a minimum LPC concentration for fusion was not identified in that work). We have ignored the kinetics of partitioning between soluble and membrane phases, but we note that the time delay of ~10 s between LPC addition and complete fusion in the experiments of Chernomordik et al. is similar in magnitude to the characteristic time for LPC phase partitioning measured by Needham and Zhelev.

### Table Captions

Table S1- Spontaneous curvature of lipid species considered here. The spontaneous curvature of ergosterol has not been measured to the best of our knowledge. Hence we used the value measured for its analog in animal species, cholesterol.

| ***Lipid Species*** | ***Spontaneous curvature, $C_i^o$ $(\mathbf{nm^{-1}})$*** | ***Reference*** |
|---|---|---|
| DOPS, No $Ca^{2+}$ | 0.069 | (18) |
| DOPS, with $Ca^{2+}$ | -0.15 | (19) |
| DOPC | -0.11 | (20) |
| DOPE | -0.35 | (21) |
| LPC | 0.26 | (16) |
| Asolectin | -0.13 | (14) |
| Ergosterol | -0.4 | (20) |

#### Figure Captions

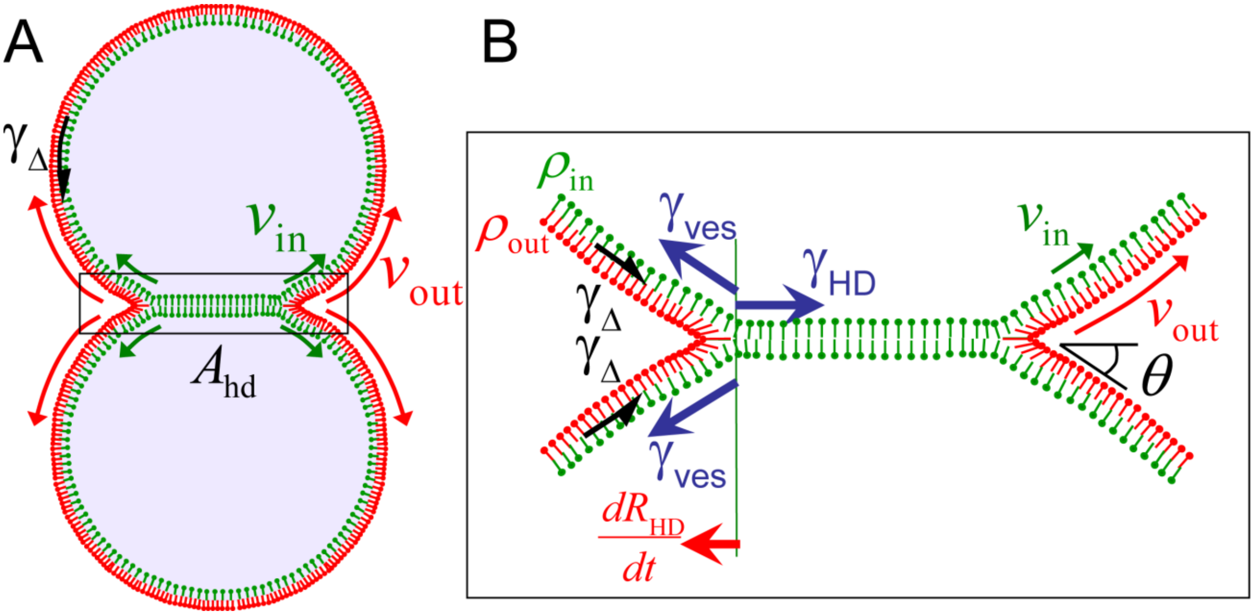


Figure S1- Growth of hemifusion diaphragm. (A) A pair of hemifused vesicles with HD area $A_{HD}$, and total vesicle surface area $A_{ves}$. (B) Blow up of the HD region. HD tension, $\gamma_{HD}(t)$ and vesicle tension $\gamma_{ves}(t)$ evolve with time according to eqs. (1)-(4). As the HD grows, the lipid density in the outer leaflet ($\rho_{out}$, red lipids) increases as lipids are forced out of the HD region and compressed. This is the fundamental driving force for HD growth, due to vesicle tension $\gamma_{ves}(t)$ and cation condensation effects. However, the lipid density of inner leaflet ($\rho_{out}$, green lipids) is lower than the outer leaflet, leading to an interleaflet tension $\gamma_\Delta$ which depends on the local areal strain $\Delta\rho/\rho_0$, eq. (1). (Adapted from J. M. Warner, B. O'Shaughnessy, Evolution of the hemifused intermediate on the pathway to membrane fusion. *Biophys J* **103**, 689-701 (2012))

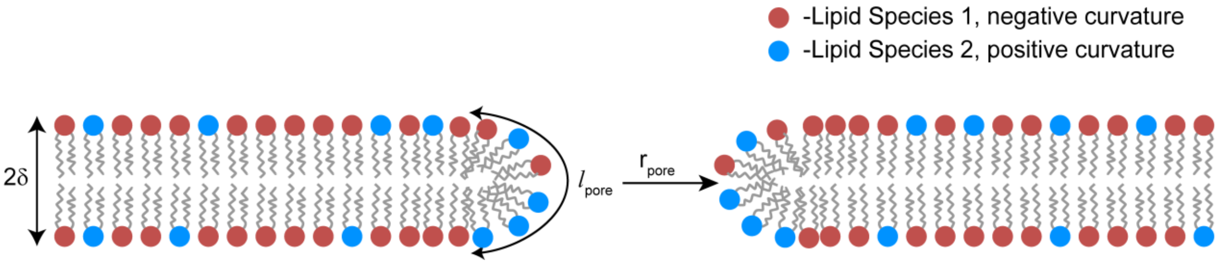


Figure S2- Schematic of a nascent pore composed of two lipid species (red and blue heads). The bilayer has thickness $2\delta$ and radius $r_{\mathrm{pore}}$. The longitudinal length of the pore is $l_{\mathrm{pore}}$. If lipid species 1 is pore-loving (pore-hating), it will partition into the pore (planar bilayer), lowering the line tension $\tau$ (eq. S20 and S26) and reducing the barrier to nascent pore dilation $\gamma^{\mathrm{crit}} = \pi\tau/\mathrm{kT}$ (*Flickering pore kinetics (HD⇌ HD**, eqs. S12, S13)).

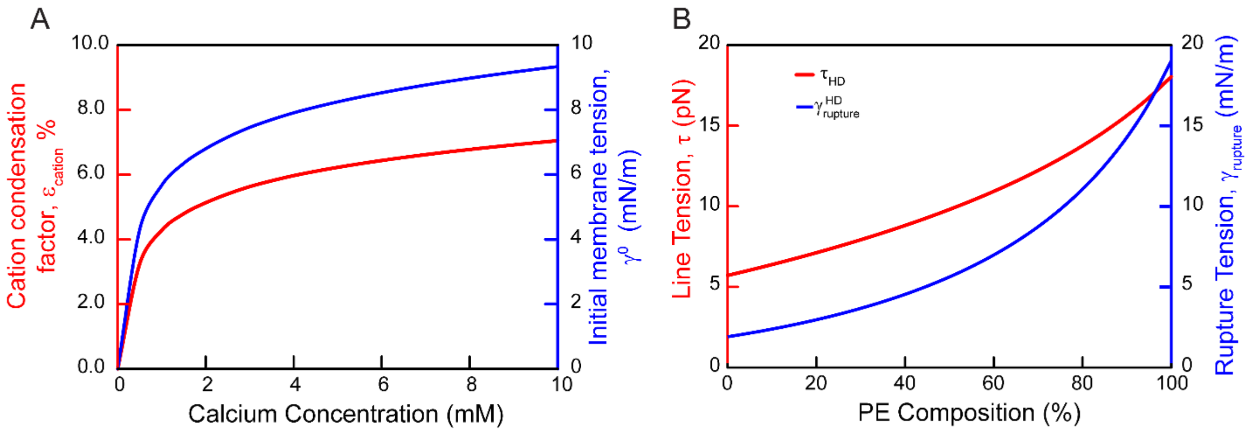


Figure S3- Calcium and PE affect membrane tension and susceptibility to rupture. (A) Calcium concentration increases both the condensation factor and the initial membrane tension. Calcium concentration controls the cation condensation factor, fit from data of refs (10, 22) for pure lipid species and taking the composition weighted average as described in *Estimation of divalent cation-induced membrane tension preceding adhesion* above. $\epsilon_{\mathrm{cation}}$, combined with the adhesion energy $W$ to set the initial membrane tension $\gamma^0$. (B) Increasing PE increases pore line tension and the membrane rupture threshold. Line tension increases with negative spontaneous curvature of the lipid components of the bilayer (eq. S21 & S27). We calculate the rupture tension $\gamma_{\mathrm{rupture}}$ by setting the pore growth rate $k_{\mathrm{grow}}$ equal to the pore resealing rate $k_{\mathrm{seal}}$. $k_{\mathrm{grow}}$ decreases as line tension increases (eq. (5) of main text) For clarity we only show $\tau_{\mathrm{HD}}$ and $\gamma_{\mathrm{rupture}}^{\mathrm{HD}}$, though the behavior for the vesicle surface is qualitatively similar.

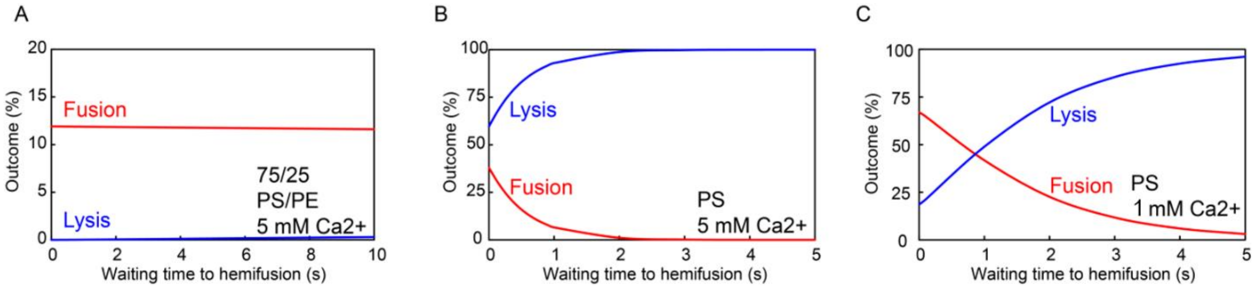


Figure S4-Waiting time for hemifusion. For the results presented in the main text it was assumed that the adhesion-to-hemifusion transition (eq. S12, S13) is instantaneous. We solved the model equations using finite waiting times. (A) For the experiments of Kachar and Rand, predicted outcomes are almost entirely insensitive to the hemifusion rate because the strongly negative spontaneous curvature and correspondingly high pore line tension of the PS/PE mixture protects the vesicle surface against rupture. (B) For the experiments of Reese and Rand, predicted outcome distributions are sensitive to waiting times from 0 to ~1 s. For longer waiting times, essentially all vesicles lyse before hemifusion. That the experimentally determined outcome distribution closely matches our predictions for very short waiting times suggests that hemifusion is in fact rapid in the PS vesicles and is not rate-limiting on the fusion pathway. Note that qualitatively, model predictions are insensitive to the waiting times, as dead-end hemifusion is essentially zero while the fusion-to-lysis ratio is just adjusted quantitatively. (C) We used the same parameters as in (B) but reduced the calcium concentration to 1 mM which lowered the vesicle tension and thus lowered the likelihood of vesicle rupture. In these conditions the outcome distribution is much less sensitive to the adhesion-to-hemifusion transition than in higher calcium concentrations.
